# A comprehensive analysis of *APOE* genotype effects on human brain structure in the UK Biobank

**DOI:** 10.1101/2023.06.19.543571

**Authors:** Verena Heise, Alison Offer, William Whiteley, Clare E Mackay, Jane M Armitage, Sarah Parish

**Affiliations:** Clinical Trial Service Unit and Epidemiological Studies Unit, Nuffield Department of Population Health, University of Oxford, Oxford, United Kingdom; Medical Research Council Population Health Research Unit, Nuffield Department of Population Health, University of Oxford, Oxford, United Kingdom; Centre for Clinical Brain Sciences, University of Edinburgh, Edinburgh, United Kingdom; Department of Psychiatry, University of Oxford, Oxford, United Kingdom; Wellcome Centre for Integrative Neuroimaging, Oxford Centre for Human Brain Activity, University of Oxford, Oxford, UK

**Keywords:** Magnetic resonance imaging, Apolipoprotein E, Alzheimer’ disease, UK Biobank, grey matter, white matter

## Abstract

Alzheimer’s disease (AD) risk is increased in carriers of the apolipoprotein E (*APOE*) ε4 allele and decreased in ε2 allele carriers compared with the ε3ε3 genotype. The aim of this study was to determine whether: *APOE* genotype affects brain grey (GM) or white matter (WM) structure; and if differences exist, the age when they become apparent and whether there are differential effects by sex. We used cross-sectional magnetic resonance imaging data from ~43,000 (28,494 after pre-processing) white British cognitively healthy participants (7,446 *APOE* ε4 carriers) aged 45-80 years from the UK Biobank cohort and investigated image-derived phenotypes (IDPs). We observed no statistically significant effects of *APOE* genotype on GM structure volumes or median T2* in subcortical structures, a measure related to iron content. Volume of white matter hyperintensities differed significantly between *APOE* genotype groups with higher volumes in *APOE* ε4ε4 (effect size 0.14 standard deviations [SD]) and ε3ε4 carriers (effect size 0.04 SD) but no differences in ε2 carriers compared with ε3ε3 carriers. WM integrity measures in the dorsal (mean diffusivity [MD]) and ventral cingulum (MD and intracellular volume fraction), posterior thalamic radiation (MD and isotropic volume fraction) and sagittal stratum (MD) indicated lower integrity in *APOE* ε4ε4 carriers (effect sizes around 0.2-0.3 SD) and ε3ε4 (effect sizes around 0.05 SD) carriers but no differences in ε2 carriers compared with the *APOE* ε3ε3 genotype. Effects did not differ between men and women. *APOE* ε4 homozygotes appeared to have lower WM integrity specifically at older ages with a potentially steeper decline of WM integrity from the age of 60 that corresponds to around 5 years greater “brain age”. *APOE* genotype affects various white matters measures, which might be indicative of preclinical AD processes. This hypothesis can be assessed in future when clinical outcomes become available.

## Introduction

Carriers of the apolipoprotein E gene (*APOE*) ε4 allele (~ 25% of white Europeans) have a higher risk of Alzheimer’s disease (AD) (odds ratio of ~3 for heterozygotes and ~15 for homozygotes) while carriers of the rarer ε2 allele have a lower AD risk (odds ratio of 0.6) compared with carriers of the most common *APOE* ε3ε3 genotype [1, 2]. There may be an interaction between *APOE* genotype and sex because only female but not male ε4 heterozygotes showed increased AD risk [2], however this view has recently been challenged by a much larger meta-analysis [3].

Magnetic resonance imaging (MRI) can be used to identify changes in brain structure in symptomatic AD patients [4]. In studies of familial AD cases and controls, differences in cortical thickness between participants carrying autosomal dominant AD genes and non-carriers can be observed years before onset of symptoms with some regions such as the precuneus and parahippocampus showing thinning ~10 years before disease onset [5]. Therefore, changes in brain structure in healthy adults carrying *APOE* ε4 might be MRI biomarkers of pre-symptomatic AD that could identify potential participants for clinical trials of AD prevention or treatment before the onset of clinical symptoms, at which point interventions might be more effective [6].

Although small cross-sectional studies reported significant effects of *APOE* genotype on grey matter (GM) ([7], N=65 ε4 carriers) and white matter (WM) structures ([8], N=275 ε4 carriers), most studies of *APOE* genotype effects did not report significant differences ([9–14], N=50-1070 ε4 carriers). In adolescents *APOE* genotype was not associated with differences in brain structure in some ([15, 16], N=120-340 ε4 carriers), although not all studies ([17, 18], N=60-310 ε4 carriers). However, these studies were likely underpowered because they were small, or aggregated data from different MRI scanners with different data acquisition protocols and data analysis methods.

To more reliably estimate differences by *APOE* genotype, studies of a larger number of participants scanned with identical MRI hardware and data acquisition protocols are needed. In this study, we assessed whether, in about 43,000 UK Biobank participants with MRI brain scanning with a common protocol, *APOE* genotype affects brain grey or white matter structure and if differences exist, at what age they become apparent and whether there are differential effects by sex.

## Materials and Methods

### Study design and participants

The UK Biobank (UKB) cohort is a large prospective epidemiological study of around 0.5 million participants [19]. Between 2006 and 2010 men and women aged 40 to 69 years were recruited and data collected on their lifestyle, environment, medical history and physical measures, along with biological samples. In 2014, UKB started to collect brain, heart and body imaging data from 100,000 UKB participants [20]. Data from 43,796 UK Biobank participants with pre-processed neuroimaging data released in February 2021 were included here.

### Ethical approval

UKB has ethics approval from the North West Multi-centre Research Ethics Committee (MREC), which covers the UK (for more information, see: http://www.ukbiobank.ac.uk/ethics/). This project was approved by UK Biobank (8835).

### Genetic data

Procedures for genotyping in the UK Biobank are described in more detail elsewhere [21] and online on http://www.ukbiobank.ac.uk/scientists-3/genetic-data. For this project, *APOE* genotype information was derived from the combined allelic information of two SNPs, rs423958 and rs7412. Imputed and directly genotyped SNPs were identical where both were present but more participants had complete imputed genetic information available (99.8% vs ~85% for direct genotyping), so imputed genotypes were used in all analyses.

### Neuroimaging data

The neuroimaging data acquisition and pre-processing pipelines have been described previously [22, 23]. In brief, data used in this project were acquired at three imaging centres using identical MRI scanners (3T Siemens Skyra, software VD13) and the standard Siemens 32-channel receive head coil. The scanning protocol consisted of structural T1-weighted MRI (T1), resting-state functional MRI (rsfMRI), task fMRI (tfMRI), T2-weighted fluid-attenuated inversion recovery (FLAIR) imaging, diffusion MRI (dMRI) and susceptibility-weighted MRI (swMRI). These different modalities provide information on GM structure (T1), functional connectivity (rsfMRI), brain activation in response to a specific task (tfMRI), volumes of white matter hyperintensities (FLAIR), WM structure/ connectivity (dMRI) and vascular lesions/ iron content (swMRI). Automated pre-processing pipelines were developed for the UK Biobank imaging data to create image-derived phenotypes (IDPs). For T1-weighted data, the IDPs analysed in this study were: total GM, WM and cerebrospinal fluid (CSF) volumes and GM volumes for cortical regions of interest (ROIs), not including the cerebellum, derived using the FAST algorithm as well as subcortical GM volumes derived using the algorithm FIRST. For FLAIR imaging, total volume of white matter hyperintensities (WMHs) was provided as an IDP. All volumetric IDPs (GM and WMH volumes) were corrected for head size using the volumetric scaling factor provided in the UK Biobank data before running statistical tests. The IDPs used from diffusion MRI were fractional anisotropy (FA), mean diffusivity (MD) and three neurite orientation dispersion and density imaging (NODDI) measures (intra-cellular volume fraction [ICVF], isotropic/ free water volume fraction [ISOVF] and orientation dispersion [OD]) averaged across skeletonised white matter tracts, excluding tracts in the cerebellum. Note that the two IDPs indicating different parts of the cingulum were renamed in this study to use more “standard” notation: the cingulum cingulate gyrus corresponds to the dorsal cingulum in standard notation and the cingulum hippocampus to the ventral cingulum [24]. For swMRI IDPs that denoted median T2* across subcortical structures were investigated. Functional data were not investigated in this study, only 1 IDP that indicated motion during the resting-state fMRI scan was used as a confounder in later analyses.

### Medical history and physical measures

Self-reported medical history was used to define history of neurological or cerebrovascular disease or neurological cancers. Additionally, cardiovascular disease (CVD: angina, myocardial infarction, heart or cardiac problem, peripheral vascular disease, leg claudication/ intermittent claudication, arterial embolism, aortic aneurysm, aortic aneurysm rupture), CVD risk factors (hypertension, high cholesterol or diabetes, no distinction between type 1 and type 2), treated hypertension, hypocholesteraemia, or diabetes as well as maternal and paternal AD/ dementia family history were defined by participant report. Participants were classified as hypertensive if the average of two blood pressure measurements at the imaging visit were >= 140 mmHg for systolic blood pressure and/ or >= 90 mmHg for diastolic blood pressure.

Measurements of standing height and weight at the imaging visit were used to calculate body-mass index (BMI) for participants. If standing height was not measured at the imaging visit, measurements from previous visits were used but if weight was not measured at the imaging visit BMI was not calculated and treated as missing.

### Other phenotype data

Townsend deprivation index was used as a measure of socioeconomic status [25]. Education was categorised into: no qualification, O-levels or equivalent, A levels or equivalent and University degree or equivalent (includes vocational and professional qualifications).

### Exclusion criteria

Participants were excluded: (i) if they reported medical history of conditions that might lead to structural brain abnormalities or cognitive impairment, i.e. any central or peripheral nervous system or nerve tumour, any chronic neurological illness or nervous system trauma, any cerebrovascular disease or intracranial haemorrhage, any infection of nervous system, cranial nerve palsy, spinal cord disorder, epilepsy, or cerebral palsy, (ii) if their UK Biobank genetic quality control flag indicated unusually high heterozygosity or >5% missing genotype rate, or if information on kinship indicated that participants were related to individuals within the sample (3rd degree or closer), or if participants did not have a Caucasian genetic ethnic background, or if there was a mismatch between self-reported sex and genetic sex, or if they carried *APOE* genotypes other than ε2/ε2, ε2/ε3, ε3/ε3, ε3/ε4 or ε4/ε4, *(iii) if* they did not have complete MRI datasets with data from all structural modalities and the rsfMRI scan, were outliers (3 standard deviations (SD) above the mean) on head motion during the rsfMRI acquisition or on the number of outlier slices detected during pre-processing of the dMRI data. For the computation of cut-offs, all available MRI data were used irrespective of scan site.

### Statistical analyses

Data were analysed with SAS version 9.4 and figures were prepared using R version 3.6.2 (https://cran.r-project.org/). *APOE* genotype groups were classified as ε2 carriers (genotypes ε2ε2 and ε2ε3), ε3 homozygotes (ε3ε3), ε4 heterozygotes (ε3ε4) or ε4 homozygotes (ε4ε4).

Baseline characteristics were compared between *APOE* genotype groups with analysis of variance (ANOVA) for continuous variables (age, Townsend deprivation index, BMI) or Pearson’s χ^2^ test for categorical variables (sex, education, CVD, hypertension, high cholesterol, diabetes, dementia family history).

To analyse effects of *APOE* genotype on brain structure, multiple linear regression was run for every IDP of interest to determine whether standardised means differed significantly between *APOE* genotype groups. The IDP was used as the dependent variable and *APOE* genotype group and confounders as independent variables. One IDP (volume of WMHs) was log-transformed before statistical analyses because the distribution was positively skewed. Before analyses all IDPs were standardised by dividing by their standard deviation. Results were displayed in ‘Manhattan’ style plots of p-values for the heterogeneity of each IDP with *APOE* genotype categories.

#### Confounders

Models included age and age^2^, sex, age*sex, age^2^*sex, educational attainment and twenty principal components of genetic ancestry provided by UK Biobank. Models included neuroimaging confounders, as described previously [26, 27]: the volumetric scaling factor to correct for head size (only used in analyses of non-volumetric IDPs, volumetric IDPs were corrected for head size before statistical analyses), head motion and head motion^2^ from the rsfMRI analysis and head position (x, y, z) in the MRI scanner as linear and quadratic terms. Since z position and table position in the MRI scanner are highly anticorrelated (r = −0.91), table position was not used as confounder. Month of scan (calculated from the date of first scan) was also added as a categorical confounder to account for temporal drifts in the data. These confounders roughly correspond to the “simple” set of confounders in [27]. Analyses where data from different scan sites were combined (e.g. the replication dataset) were additionally adjusted for the scanner site and the interaction between the site and each imaging confounder.

#### Correcting for multiple comparisons

Bonferroni correction was applied to correct for multiple comparisons as follows. The alpha level of 0.05 was divided by the total number of modalities investigated, i.e. 4 (T1, dMRI, swMRI and FLAIR), to derive one alpha level per modality, i.e. 0.0125. Within modality, this alpha level was then divided by the number of brain regions investigated: 113 for T1, 200 for dMRI (40 regions*5 different diffusion measures), 14 for swMRI and 1 for FLAIR. This resulted in the following Bonferroni correction thresholds for the different modalities: p < 1*10^-4^ for T1, p < 6*10^-5^ for dMRI, p < 9*10^-4^ for swMRI and 0.0125 for FLAIR.

#### Discovery and replication dataset

The data were split based on the MRI scanner used for data acquisition. Cheadle was the discovery dataset (n=17239) and Newcastle (n=7465) and Reading (n=3790) were combined as the replication dataset to determine whether IDPs passing the level of significance in the discovery analysis (‘hits’) could be replicated. The p-value threshold for significance at replication per modality was 0.0125 divided by the number of significant hits in that modality in the discovery cohort.

#### Post-hoc analyses of hits in the discovery cohort

For the replicating hits within each modality likelihood ratio tests were used to assess whether hits in left and right hemisphere IDPs remained statistically significant after adjustment for their average level, and if they did not then the average level was used in further analyses.

Effects (in units of standard deviation of the IDP) of each *APOE* genotype in comparison to the ε3ε3 genotype were estimated. Analyses were first run separately for the discovery and replication dataset and then data from all scan sites were combined in one analysis, adjusted for site.

Sensitivity analyses were conducted altering the level of adjustment for age and neuroimaging confounders: finer adjustment for age using individual years of age as a single year categorical variable, no adjustment for neuroimaging confounders, rank inverse-normal transformation of all IDPs, age and neuroimaging confounders except for imaging centre. All sensitivity analyses were adjusted for twenty principal components of genetic ancestry.

#### Subgroup analyses

Subgroup analyses were conducted in a combined analysis of all datasets for hits that achieved replication. Differences in the strength of associations by sex were investigated in *APOE* ε3ε4 and ε4ε4 carriers compared with the *APOE* ε3ε3 genotype group. To determine at what age *APOE* genotype differences become apparent, two subgroup analyses were conducted: first, participants were divided into two age groups age (<65 versus ≥65 years) and differences in the strength of associations were investigated for those two subgroups in *APOE* ε3ε4 and ε4ε4 carriers compared with the *APOE* ε3ε3 genotype group. Second, a more fine-grained analysis was conducted with participants divided into five age groups (<55, 55-59, 60-64, 65-69, ≥70 years) and strength of associations were compared between all *APOE* genotype groups in each of these age bins. Adjusted rate of cross-sectional change with age at imaging was calculated for all *APOE* genotype groups from linear regression of IDPs on age at imaging, principal components of genetic ancestry and the other imaging confounders.

### Pre-registration

The pre-registered study protocol is available on doi: <u>10.17605/OSF.IO/BRN3H</u>. Changes from the pre-registered protocol are described in the supplementary methods.

### Open data

UK Biobank is an open access resource and researchers can apply to use the data following procedures outlined on their website: https://www.ukbiobank.ac.uk/enable-your-research.

## Results

After data cleaning steps and applying exclusion criteria, these analyses include 28,494 participants (Supplementary Figure 1).

### Participant characteristics at the imaging visit

With absence of an *APOE* ε2 allele or a greater number of ε4 alleles, people had a greater prevalence of hypercholesterolemia and family history of AD (Table 1). *APOE* ε4 carriers were a median 1 year younger and a had a lower BMI compared with other genotypes. Other characteristics were similar by genotype.

**Table 1.** Participant characteristics at the imaging visit by *APOE* genotype. P values refer to comparisons using χ^2^ tests for categorical variables and ANOVAs for continuous variables.

|  | ε2 carriers<br>(n=3842) | ε3ε3<br>(n=17 206) | ε3ε4<br>(n=6794) | ε4ε4<br>(n=652) | P value |
| --- | --- | --- | --- | --- | --- |
| Age (years) |  |  |  |  |  |
| Mean (SD) | 64.1 (7.7) | 64.1 (7.6) | 63.6 (7.6) | 63.2 (7.3) | 3.0×10<sup>-6</sup> |
| Median [Min,Max] | 65.0 [46.0,81.0] | 65.0 [45.0,82.0] | 64.0 [46.0,82.0] | 64.0 [47.0,79.0] |  |
| Sex |  |  |  |  |  |
| Female | 2057 (53.5%) | 9099 (52.9%) | 3659 (53.9%) | 372 (57.1%) | 0.12 |
| Townsend Deprivation Index |  |  |  |  |  |
| Mean (SD) | -2.07 (2.66) | -2.07 (2.61) | -2.01 (2.64) | -2.16 (2.47) | 0.38 |
| Median [Min,Max] | -2.77 [-6.26,9.23] | -2.73 [-6.26,9.74] | -2.68 [-6.26,8.91] | -2.84 [-6.26,7.18] |  |
| Education |  |  |  |  |  |
| No qualifications | 208 (5.4%) | 922 (5.4%) | 339 (5.0%) | 29 (4.4%) | 0.90 |
| O-levels/CSE/equivalent | 443 (11.5%) | 1976 (11.5%) | 773 (11.4%) | 80 (12.3%) |  |
| A-levels | 183 (4.8%) | 840 (4.9%) | 335 (4.9%) | 27 (4.1%) |  |
| Degree/professional | 3007 (78.3%) | 13467 (78.3%) | 5347 (78.7%) | 516 (79.1%) |  |
| Body Mass Index (kg/m<sup>2</sup>) |  |  |  |  |  |
| Mean (SD) | 26.4 (4.2) | 26.3 (4.2) | 26.3 (4.3) | 25.7 (3.9) | 1.0×10<sup>-3</sup> |
| Median [Min,Max] | 25.8 [15.7,46.9] | 25.8 [13.4,57.2] | 25.7 [14.1,53.4] | 25.3 [16.0,42.3] |  |
| AD family history |  |  |  |  |  |
| Mother | 550 (14.3%) | 2768 (16.1%) | 1538 (22.6%) | 189 (29.0%) | 6.2×10<sup>-50</sup> |
| Father | 305 (7.9%) | 1450 (8.4%) | 860 (12.7%) | 109 (16.7%) | 1.4×10<sup>-32</sup> |
| Medical Conditions |  |  |  |  |  |
| Hypertension | 1958 (51.0%) | 8793 (51.1%) | 3432 (50.5%) | 318 (48.8%) | 0.60 |
| Diabetes | 106 (2.8%) | 443 (2.6%) | 154 (2.3%) | 8 (1.2%) | 0.063 |
| Hypercholesterolemia | 672 (17.5%) | 3978 (23.1%) | 1786 (26.3%) | 200 (30.7%) | 2.8×10<sup>-27</sup> |
| CVD | 30 (0.8%) | 157 (0.9%) | 80 (1.2%) | 2 (0.3%) | 0.045 |

### Discovery analyses

In the discovery cohort, WM integrity of four tracts showed statistically significant differences between *APOE* genotype groups: dorsal cingulum, ventral cingulum, posterior thalamic radiation, and sagittal stratum (Figure 1a). 3D-images of those brain structures can be found on https://identifiers.org/neurovault.collection:9357.

**Figure 1.**
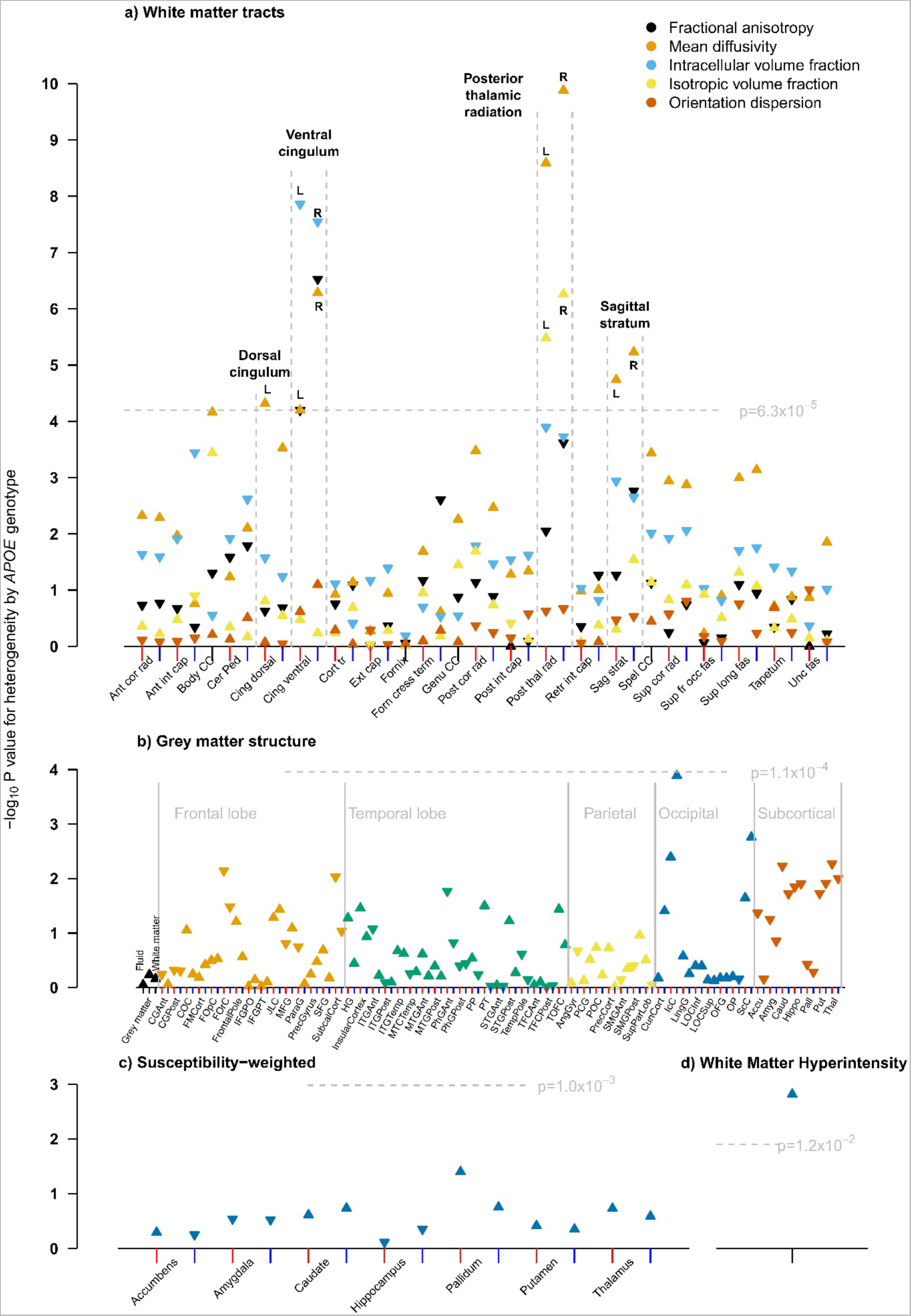
Manhattan plots showing p-values for heterogeneity across *APOE* genotypes from multiple linear regression analyses in the discovery cohort to determine *APOE* genotype differences on different measures of brain structure: white matter (panel a), grey matter (panel b), swMRI (panel c) and volume of white matter hyperintensities (panel d). For IDPs that have left and right components, the x positions of left and right components are indicated with red and blue tick marks respectively. For grey matter structure, the markers have been ordered by brain region: whole brain, frontal lobe, temporal lobe parietal, occipital, and subcortical. An upwards pointing triangle indicates that the IDP is higher in participants with the *APOE* ε4ε4 genotype compared with the ε3ε3 genotype, a downwards pointing triangle indicates it is lower.

The WM integrity measures that showed genotype differences were: MD (left only) for the dorsal cingulum; ICVF, MD (both left and right) and FA (right only) for the ventral cingulum; MD and ISOVF (both left and right) for the posterior thalamic radiation; and MD (left and right) for the sagittal stratum. In each tract, WM integrity was lower (increased MD and ISOVF, decreased FA and ICVF) in *APOE* ε4ε4 carriers compared with the ε3ε3 genotype. Additionally, total WMH volume differed statistically significantly between *APOE* genotype groups with higher volumes in carriers of *APOE* ε4ε4 genotype (Figure 1d).

In contrast, there were no statistically significant differences in volumes of grey matter structures (Figure 1b) or median T2* in subcortical structures (Figure 1c) between *APOE* genotype groups.

### Replication analyses

The significance threshold for the replication analysis with data from the other two imaging centres was 0.0125 / 12 hits = 1.0*10^-3^ for the analysis of white matter IDPs and 0.0125 for the volume of white matter hyperintensities.

The following WM integrity IDPs differed significantly by genotype (in the same direction) in both the discovery and replication cohort: MD in the left dorsal cingulum; ICVF and MD in the left and right ventral cingulum; MD and ISOVF in the left and right posterior thalamic radiation; and MD in the left and right sagittal stratum (Figure 2a). The association of *APOE* genotype with WMH volume was also replicated (Figure 2b). An analysis of all IDPs in the replication cohort provided visual confirmation that patterns in the Manhattan plots for the discovery cohort were replicated (Supplementary Figure 2).

**Figure 2.**
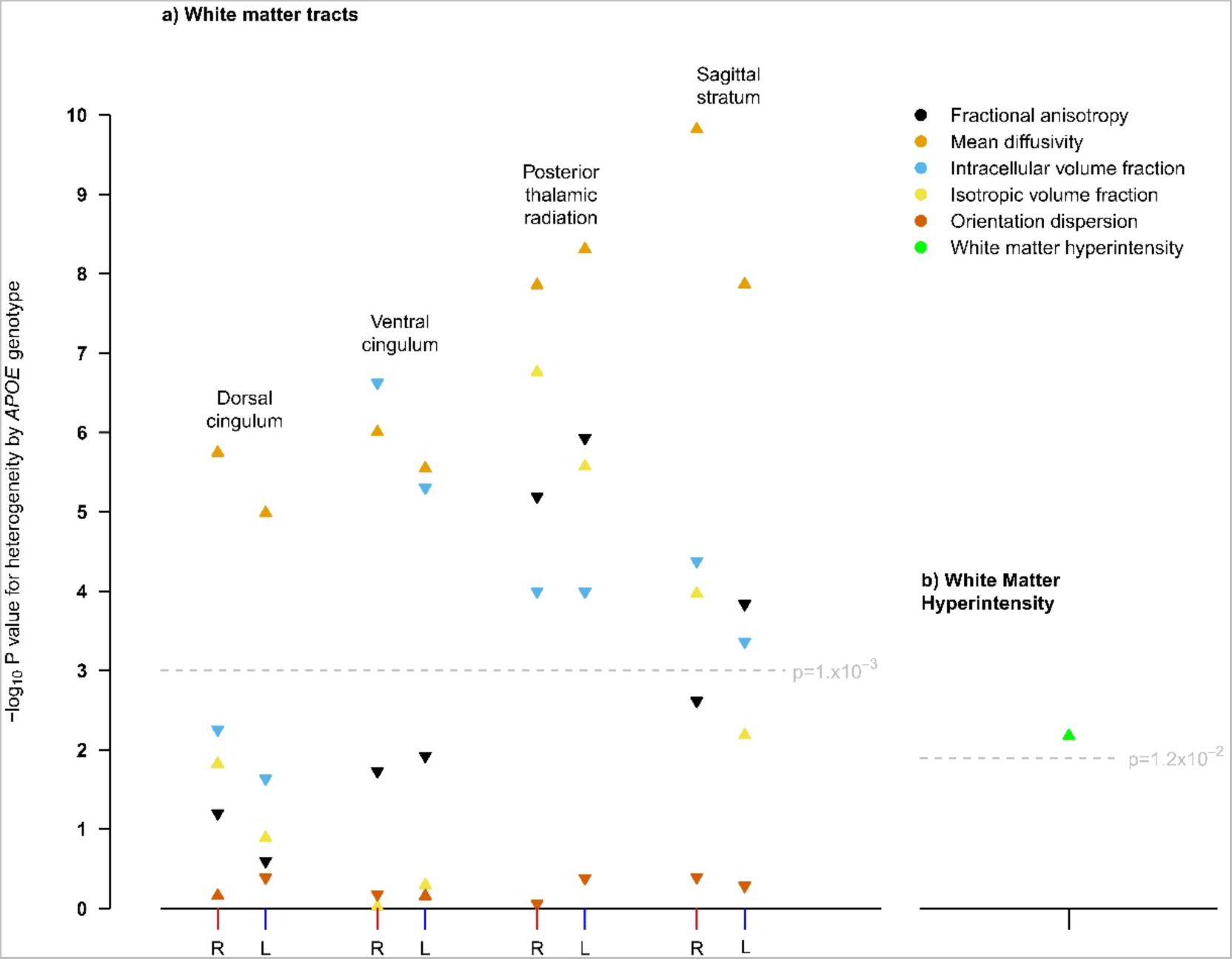
Manhattan plots showing p-values for heterogeneity across *APOE* genotypes in the replication cohort for IDPs that reached significance in the discovery cohort, i.e. white matter IDPs (panel a) and volume of white matter hyperintensities (panel b). L and R is used to distinguish left from right. An upwards pointing triangle indicates that the IDP is higher in participants with the *APOE* ε4ε4 genotype compared with the ε3ε3 genotype, a downwards pointing triangle indicates it is lower.

### Independence of left and right WM hits

In the discovery data, there was no evidence that *APOE* genotype associations with WM integrity differed by side of tract (left or right) (Supplementary Table 1), so an average of both tracts was used in subsequent analyses.

### Sensitivity analyses

Sensitivity analyses were conducted for one white matter IDP (MD in the posterior thalamic radiation). They showed very similar results for all analyses irrespective of the types of adjustments (Supplementary Figure 3).

### Comparison of WM integrity and WMH volumes between *APOE* genotype groups

For the IDPs that achieved replication for statistically significant effects of *APOE* genotype, effects were investigated further to determine, which *APOE* genotype groups differed significantly from *APOE* ε3ε3 carriers. *APOE* ε4ε4 carriers showed the largest differences of 0.14-0.31 standard deviations (SDs) compared with ε3ε3 carriers, ε3ε4 carriers showed very small but statistically significant differences of around 0.05 SDs and ε2 carriers did not differ significantly from ε3ε3 carriers. Thus, the effect size for *APOE* ε4 homozygotes was much stronger than twice the effect for *APOE* ε3ε4 carriers (Figure 3).

**Figure 3.**
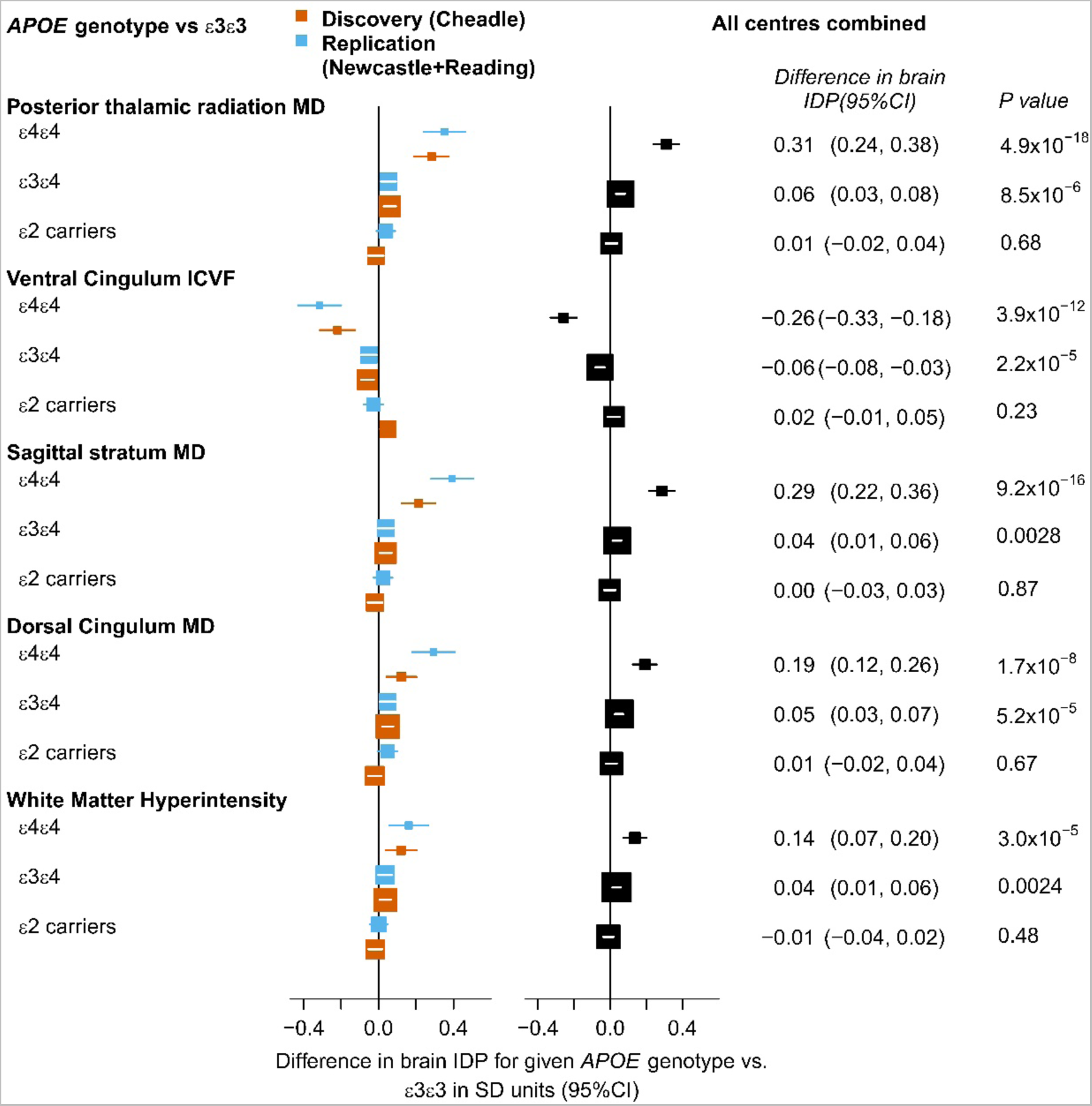
Differences in WM integrity and WMH volume between *APOE* genotype groups for IDPs that achieved replication for heterogeneity with *APOE* genotype. The *APOE* ε3ε3 genotype is the reference group. On the left, there are results for separate analyses for the discovery (red) and replication (blue) dataset. On the right there are results for the combined analysis of all data.

### Subgroup analyses by sex and age

For the IDPs that achieved replication for heterogeneity across *APOE* genotype, there was no strong evidence for effect modification by sex given the number of comparisons when comparing *APOE* ε3ε4 or ε4ε4 carriers with the reference genotype *APOE* ε3ε3 (Figure 4). The comparison of results of two age groups (< 65 years) and (≥ 65 years) indicated that effects differed markedly in *APOE* ε4ε4 carriers where the older age group showed double or more the effect sizes of the younger group, in particular for WM integrity in the posterior thalamic radiation and sagittal stratum; effects of the *APOE* ε3ε4 genotype were similar for both age groups (Figure 4).

**Figure 4.**
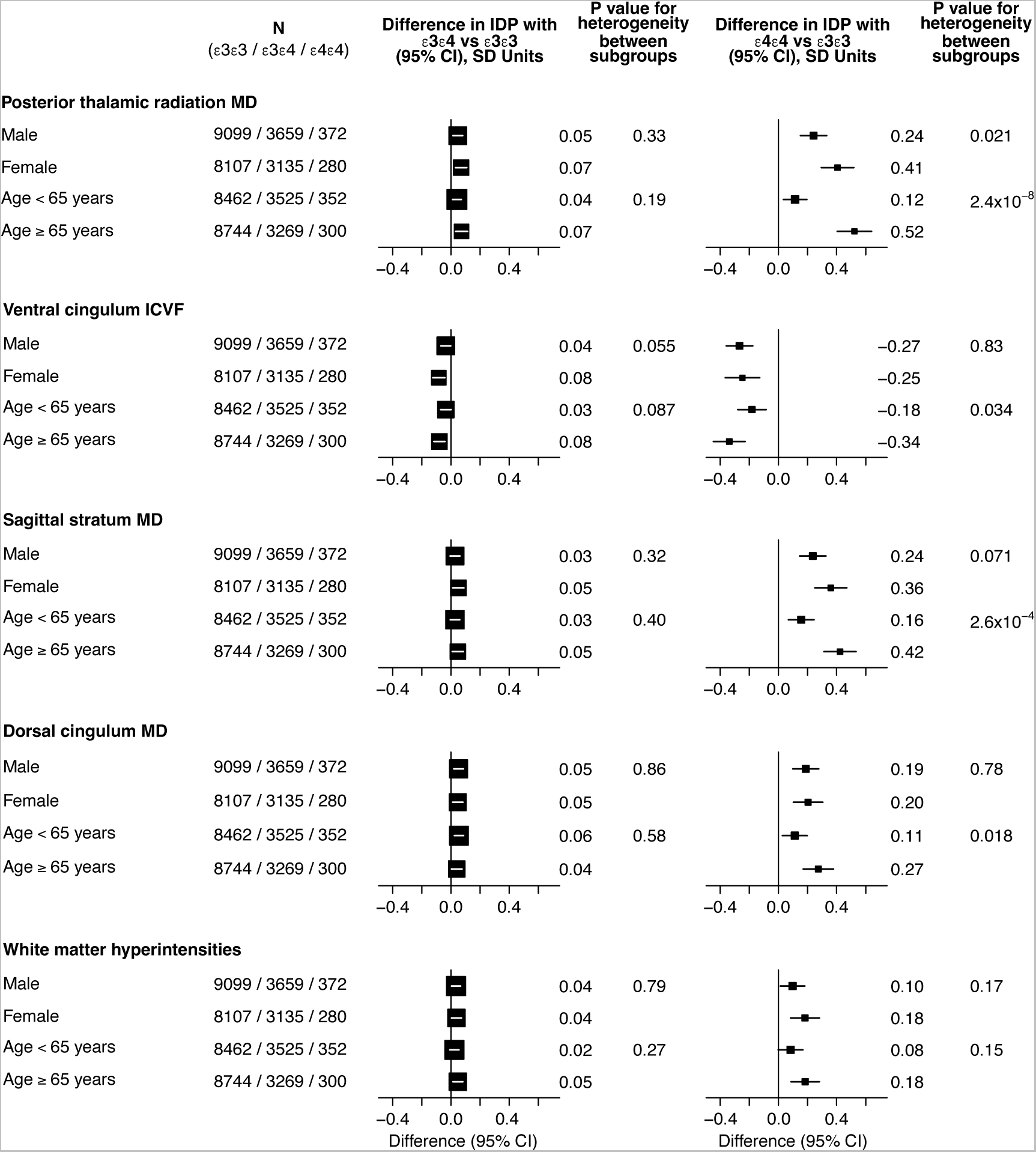
Combined analysis of all centres with subgroup analyses for the comparison of *APOE* ε3ε4 (left) and *APOE* ε4ε4 carriers (right) with the reference group *APOE* ε3ε3 by sex (male vs female) and by age group (< 65 vs ≥ 65 years) for the IDPs that achieved replication for heterogeneity by *APOE* genotype.

A more detailed analysis of 5-year age groups showed worsening WM integrity (increase in MD and decrease in ICVF) and increase in WMH volume with age for all *APOE* genotype groups in this cross-sectional analysis. For *APOE* ε2 carriers, ε3ε3 and ε3ε4 genotypes the WM integrity worsened at a similar rate with age, whereas for the *APOE* ε4ε4 genotype there was a higher rate of change per year from the age group 55-59 years. The mean difference in WM integrity between *APOE* ε4ε4 carriers and all other genotype groups from age group 65-69 years onwards corresponded to the mean IDP difference seen with around 5 years greater age cross-sectionally over the whole population (Figure 5).

**Figure 5.**
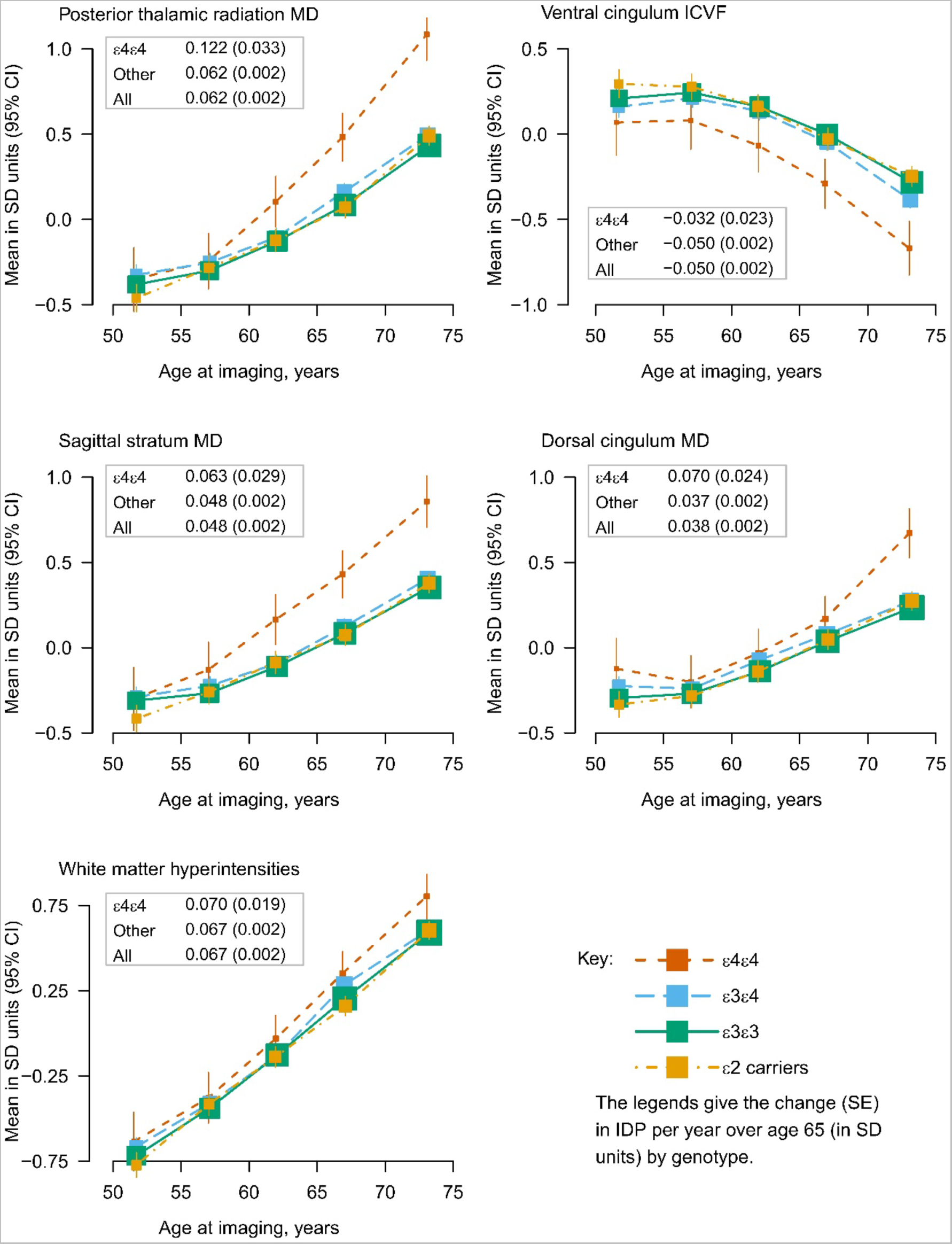
Mean levels of IDPs that achieved replication by *APOE* genotype group averaged across 5-year age groups. IDPs are standardised to a mean of zero and a standard deviation of one. *APOE* ε2 carriers are shown in yellow, ε3 homozygotes in green, ε4 heterozygotes in blue and ε4 homozygotes in red.

## Discussion

In this analysis of neuroimaging data from 28,494 UK Biobank participants we found statistically significant effects of *APOE* genotype on WM integrity and WMH volume but did not observe statistically significant effects on GM volumes or subcortical swMRI measures.

### White matter structure

Effects of *APOE* genotype on several measures of WM integrity were observed: MD in the dorsal cingulum; ICVF and MD in the ventral cingulum; MD and ISOVF in the posterior thalamic radiation and MD in the sagittal stratum. WM integrity was lower (lower ICVF, higher MD) in those with highest AD risk. However, differences were greatest between *APOE* ε4 homozygote and ε3ε3 carriers, small but consistent between ε4 heterozygotes and ε3ε3 carriers but not statistically significant between ε2 carriers and ε3ε3 carriers. Therefore, *APOE* genotype differences in WM integrity did not clearly map to AD risk, which is lowest in ε2 carriers [2], but were based on negative effects of the *APOE* ε4 allele only.

There have been conflicting findings in the literature of possible *APOE* genotype effects on white matter structure in smaller studies, e.g. positive in [28] or no associations in [9, 29]. In their analysis of a smaller UK Biobank sample Lyall et al. (2020) [30] did not find significant effects of *APOE* genotype on a general factor of FA (gFA), which is a single factor capturing common variance across all white matter tracts. We show here that the differences between genotypes are very localised and mainly driven by the *APOE* ε4ε4 genotype, which is why, in combination with the smaller study size, they might not have been picked up by Lyall et al. (2020).

The analyses of NODDI measures provide insight into the microstructural properties and biological mechanisms underlying the observed differences. ICVF is a measure of neurite density that is also related to myelination and ISOVF is thought to reflect CSF contamination [31]. Therefore, *APOE* genotype effects on WM integrity in the ventral cingulum could be driven by neuronal mechanisms and in particular reduced myelination, a process that might play a role in early AD development [32, 33]. The ventral cingulum connects the hippocampus, parahippocampal areas and entorhinal cortex with the posterior cingulate cortex and retrosplenial cortex [24], i.e. areas that show neuronal and synapse loss early in AD [34]. Reduced white matter integrity of this tract has been shown in patients with mild cognitive impairment and AD [24]. A study combining dMRI and histopathology showed that reduced FA and increased MD of the ventral cingulum were associated with higher Braak stages of neurofibrillary tangle pathology in AD patients [35]. Therefore, the findings of reduced integrity of the ventral cingulum here might be indicative of preclinical AD pathology in those at higher genetic AD risk.

In contrast, *APOE* genotype effects on MD and ISOVF of the posterior thalamic radiation could reflect localised expansion of ventricles, a well-known feature of AD pathology that correlates with cognitive decline [36]. *APOE* genotype differences in MD in the dorsal cingulum and sagittal stratum were not accompanied by differences in NODDI measures, so the biological mechanisms underlying differences in those tracts are unclear. However, there are previous reports that have shown reduced WM integrity of these tracts in cognitively impaired patients [37, 38].

### Effects of age and sex on WM integrity

We conducted an exploratory analysis of WM integrity across the age range in this sample, which indicated that *APOE* ε4ε4 carriers seem to deviate from the other *APOE* genotype groups particularly from the age group of 55-59 years. The differences in WM integrity measures in *APOE* ε4ε4 carriers compared with all other genotype groups correspond to WM integrity measure changes over 5 years in this cross-sectional study. Therefore, they might indicate faster ageing processes in older *APOE* ε4ε4 carriers. If confirmed in longitudinal studies, this would point to a role of *APOE* genotype on changes associated with preclinical AD processes rather than effects that exist across the lifespan. This would fit with some larger studies in children and adolescents that have reported no significant effects of *APOE* genotype on brain structure at younger ages [15, 16].

We did not find any evidence of differences between men and women in *APOE* genotype effects on brain structure. This would support a recent meta-analysis that reported no sex differences in *APOE* genotype effects on AD risk [3].

### White matter hyperintensities

There have been conflicting reports in the literature investigating possible *APOE* genotype effects on WMH volume, e.g. positive association in [39], no association in [40]. Significant effects of *APOE* genotype on WMH volumes have been reported previously in a smaller sample of UK Biobank data [30] and our results are consistent with that report. WMHs are one marker of cerebral small vessel disease [41]. Therefore, *APOE* genotype might affect AD risk via effects on the vasculature in the brain.

### Effect sizes

While we reliably observed effects of *APOE* genotype on WM structure and WMH volumes, the effect sizes for differences between groups were generally modest with 0.3 SDs or less. On the one hand, this has implications for follow-up MRI studies of *APOE* genotype effects on brain structure, which need to have large enough sample sizes to be able to detect small effects. On the other hand, it is unclear whether these modest changes can be translated into MRI biomarkers that would be sensitive enough to detect preclinical changes associated with AD development on an individual basis.

### Grey matter structure

In this study we did not observe any significant effects of *APOE* genotype on volumes of GM structures. This is not necessarily surprising since most studies with larger sample sizes do not report effects on GM structure in cognitively healthy participants, with many focusing specifically on the hippocampus [9, 12–14, 42]. This is the largest and most homogenous study that investigated effects on GM structure and did not find any effects. However, it is clear from histopathological studies that GM structures, particularly in the medial temporal lobe, are affected early by neuronal loss in AD development [34]. These changes might be too localised to translate into volumetric differences that can be detected with the IDP MRI measures used in this study. Additionally, histopathological studies have shown that synaptic loss can exceed and predate neuronal loss [34]. It is possible that WM integrity MRI measures investigated here are sensitive enough so that these preclinical changes in neuronal connectivity can be detected here.

### Susceptibility-weighted imaging

Similarly, we did not find any significant effects on median T2* in subcortical structures. Increased iron levels that affect median T2* [43], particularly in subcortical structures, have been reported in AD patients [44]. However, there is no study to date that investigated *APOE* genotype effects on median T2* levels. SWI is sensitive to iron accumulation and therefore bleeding events. A number of meta-analyses reported significant *APOE* genotype effects on volumes of cerebral microbleeds [39, 45]. Therefore, it might be more appropriate to use the T2* signal to identify localised microbleeds and investigate the volumes of these bleeds, similar to the analysis of WMHs in this paper, rather than averaging the signal across structures.

### Limitations

We only included UK Biobank participants with Caucasian genetic ethnic background, i.e. of white British ancestry, in this study. Therefore, the results might not be generalisable to other genetic ethnic backgrounds.

In this large-scale investigation of UK Biobank data we decided to focus on IDPs and did not analyse the native imaging data directly. It is currently unclear whether the use of IDPs, which are obtained by averaging measures such as T2* or FA/MD across structures or tracts, is the best way to identify changes that might happen on a smaller scale and only affect certain parts of a structure. This would become visible in analyses conducted on a voxel-by-voxel basis but not necessarily in the analysis of IDPs averaged across the whole structure.

Additionally, it is important to note that even where we did find significant differences, the effect sizes were modest. This is a cross-sectional study and we do not yet know which participants will go on to develop AD in the coming decades. Therefore, we can only speculate whether the differences between *APOE* genotypes in white matter integrity might be caused by preclinical AD processes. Participants in the UK Biobank have also been shown to be healthier than the general population [46] and we only included cognitively healthy participants in this study. Therefore, one possible reason why we mostly report null effects could be that a large number of participants will either not develop AD or develop AD relatively late in their life. This would affect our ability to capture effects of preclinical AD using *APOE* genotype as a substitute measure for AD risk.

However, evidence from cross-sectional histopathological and PET studies suggests that *APOE* genotype does affect processes that are relevant at preclinical AD, e.g. the accumulation of amyloid plaques [14, 47]. Our results suggest that these changes might only translate into modest structural changes visible using MRI modalities.

## Conclusion

In this comprehensive analysis of *APOE* genotype effects on brain structure in 28,494 white British UK Biobank participants we showed consistent significant effects on WMH volume and WM integrity of the dorsal and ventral cingulum, posterior thalamic radiation and sagittal stratum in men and women. We did not observe significant differences between *APOE* genotype groups in volumes of GM structures or median T2* in subcortical structures. *APOE* genotype effects on WM integrity in the ventral cingulum may be driven by neuronal mechanisms and reduced myelination. We saw that *APOE* ε4 homozygotes appear to have lower WM integrity at older ages with a potentially steeper decline of WM integrity from the age of 60 that corresponds to around 5 years greater “brain age”. Therefore, *APOE* genotype effects might be associated with preclinical AD. Whether this is the case may become clear in future, when these participants have been followed up for clinical outcomes and AD detected.

## CRediT authorship contribution statement

**Verena Heise:** Conceptualisation, Data Curation, Formal Analysis, Methodology, Project Administration, Software, Validation, Visualisation, Writing – Original Draft Preparation;

**Alison Offer:** Data Curation, Formal Analysis, Software, Validation, Visualisation, Writing – review and editing;

**William Whiteley:** Conceptualisation, Writing – review and editing;

**Clare Mackay:** Conceptualisation, Supervision, Writing – review and editing;

**Jane Armitage:** Conceptualisation, Supervision, Writing – review and editing;

**Sarah Parish:** Conceptualisation, Funding acquisition, Methodology, Resources, Supervision, Writing – review and editing

## Acknowledgements

This research has been conducted using the UK Biobank resource under application number 8835 (PI: Sarah Parish). We would like to thank UK Biobank participants and research team members who collected the data and administered the data release. We are especially grateful to the UK Biobank brain MRI team, in particular Stephen Smith and Fidel Alfaro-Almagro, who delivered pre-processed MRI data to UK Biobank. This work was supported by grants to the University of Oxford from the UK Medical Research Council through its funding of the MRC Population Health Research Unit (MC_ UU_00017/5). VH was supported through a Nuffield Department of Population Health Intermediate Fellowship. CEM was supported by the NIHR Oxford Health Biomedical Research Centre. For the purpose of open access, the authors have applied a Creative Commons Attribution (CC BY) licence to any Author Accepted Manuscript version arising.

