## Supplementary Information for "A comprehensive analysis of *APOE* genotype effects on human brain structure in the UK Biobank"

### Supplementary Methods

#### Pre-registration

The pre-registered study protocol is available on doi: 10.17605/OSF.IO/BRN3H. Compared with the pre-registration several changes were made: We investigated a larger sample because more data was available at this later time point (February 2021). The Bonferroni correction threshold was adjusted to account for the fact that different numbers of IDPs were available for each of the modalities and an individual threshold for each of the modalities was deemed more appropriate. We did not investigate any of the functional modalities because the resting-state functional connectivity data did not provide connectivity information of regions of interest (e.g. separate regions of the default-mode network). We did not use motion during the tfMRI scan as an exclusion criterion because there is a high correlation between motion during tfMRI and rsfMRI and more participants had data available from the rsfMRI scan. We did not run any image-based analyses due to resource limitations in how these very extensive datasets were accessible at our institution. The number of IDPs analysed per modality is slightly lower than in the pre-registration because all IDPs that covered the cerebellum were excluded since that region is only affected by AD pathology at very late disease stages [1]. For the analysis of WM integrity tensor mode was not included as a measure because it is unclear how to interpret the measure. We did not check for/ exclude potential outliers in the distribution of individual IDPs due to the large number of participants and IDPs that would have had to be checked and insufficient justification that those > 3SD above the mean would be outliers in this large study. We added a number of confounders to the analyses: principal components from the genetic analyses to correct for remaining population substructure, head position in the MRI scanner and month of scan based on reports by others [2, 3]. In addition to the pre-registered analyses, we also conducted exploratory subgroup analyses of the IDP hits that showed potential effects of *APOE* genotype to understand at what age effects of *APOE* genotype become most apparent.

### Supplementary Results

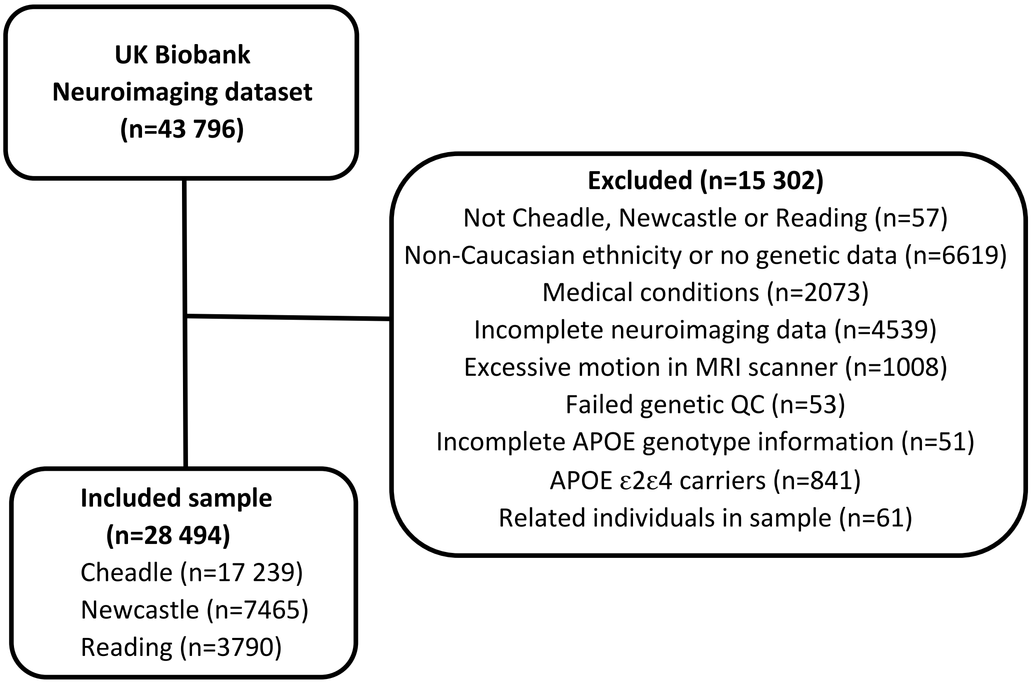


**Supplementary Figure 1.** Numbers of participants that passed each stage of exclusion criteria. Data were available for a total of 43 796 participants. After applying all exclusion criteria, data from 28 494 participants were used in the statistical analyses.

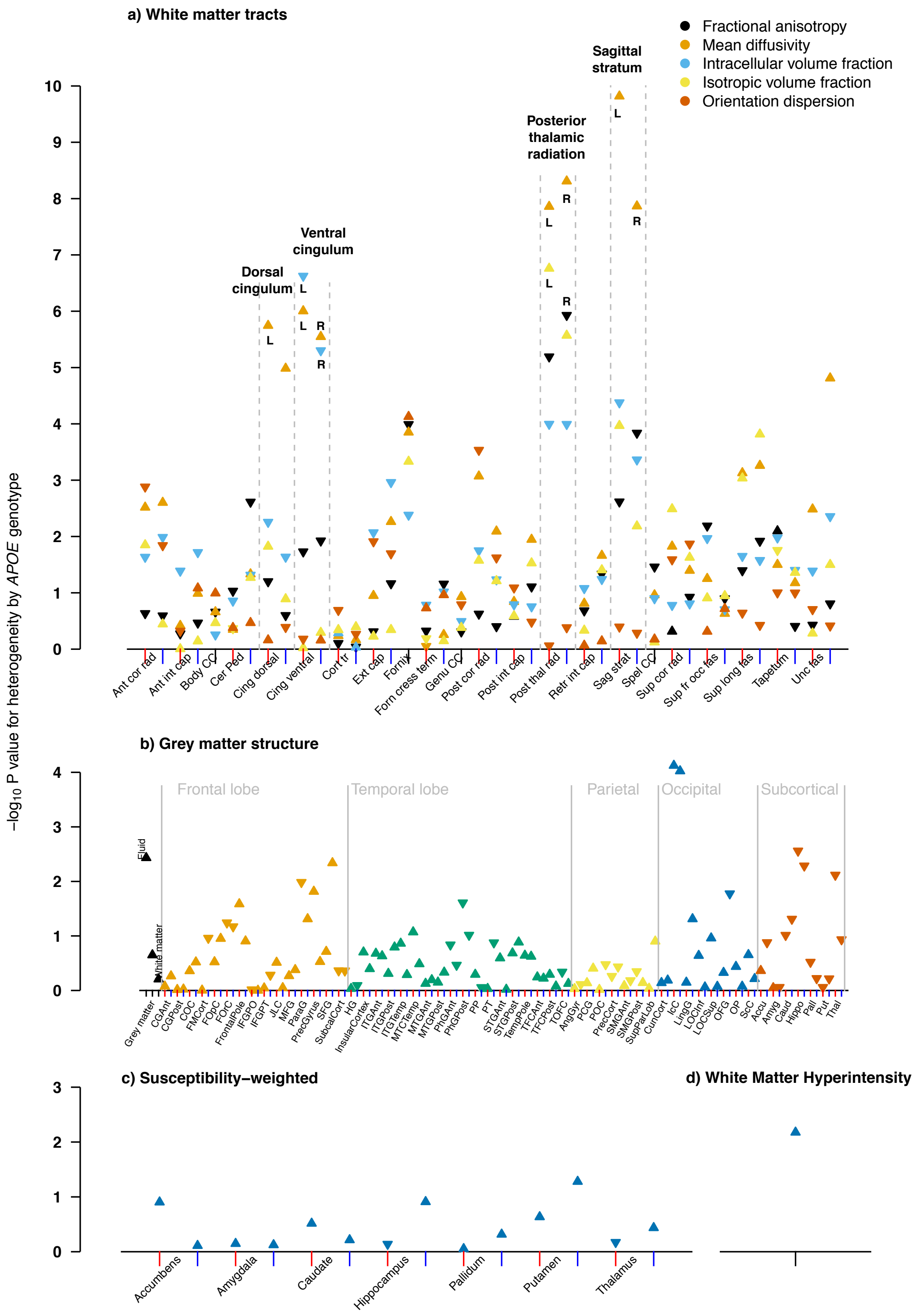


**Supplementary Figure 2.** Manhattan plots showing p-values for heterogeneity across *APOE* genotypes from multiple linear regression analyses in the replication cohort to determine *APOE* genotype differences on different measures of brain structure: white matter (panel a), grey matter (panel b), swMRI (panel c) and volume of white matter hyperintensities (panel d). For IDPs that have left and right components, the x positions of left and right components are indicated with red and blue tick marks respectively. For grey matter structure, the markers have been ordered by brain region: whole brain, frontal lobe, temporal lobe parietal, occipital, and subcortical. An upwards pointing triangle indicates that the IDP is higher in the population with the *APOE* ε4ε4 genotype compared to the population with the ε3ε3 genotype a downwards pointing triangle indicates it is lower.

**Supplementary Table 1.** Analysis of independence of left and right hits in the discovery dataset. P-values for heterogeneity of IDPs across *APOE* genotypes by hemisphere, p-values for the left (or right) hemisphere after adjustment for the right (or left) and p-values averaged over hemispheres.

| IDP | Left hemisphere | Right hemisphere | Left (or right) given average | Average over hemispheres |
|---|---|---|---|---|
| Posterior thalamic radiation MD | $2.6 * 10^{-9}$ | $1.3 * 10^{-10}$ | $6.4 * 10^{-1}$ | $1.0 * 10^{-10}$ |
| Ventral cingulum ICVF | $1.4 * 10^{-8}$ | $2.8 * 10^{-8}$ | $6.1 * 10^{-1}$ | $4.2 * 10^{-9}$ |
| Sagittal stratum MD | $1.8 * 10^{-5}$ | $5.9 * 10^{-6}$ | $9.8 * 10^{-1}$ | $2.4 * 10^{-6}$ |
| Dorsal Cingulum MD | $4.8 * 10^{-5}$ | $3.0 * 10^{-4}$ | $5.4 * 10^{-1}$ | $4.3 * 10^{-5}$ |

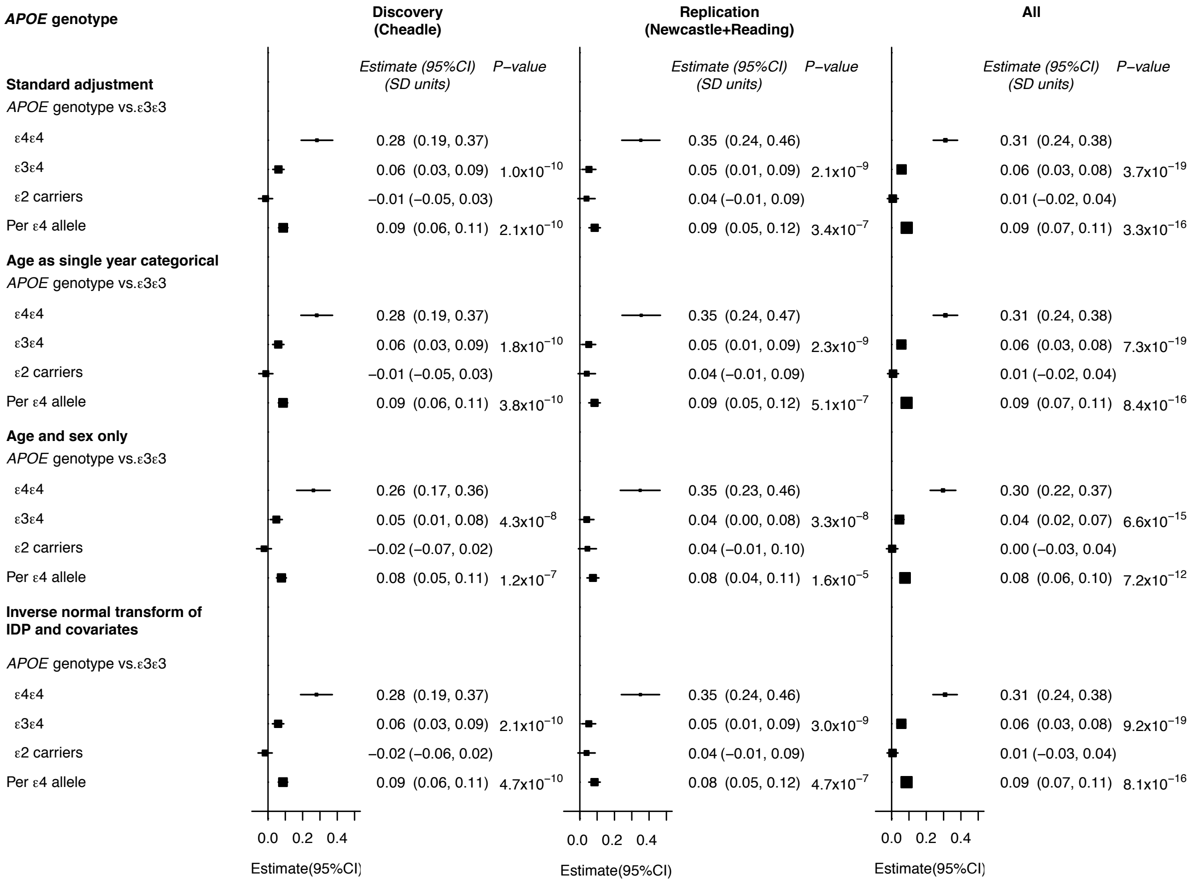


**Supplementary Figure 3.** Sensitivity analyses for MD in the posterior thalamic radiation. The top analysis uses the standard adjustment, the second one uses a finer adjustment for age with individual years of age as categorical confounder, the third one uses age and sex only with no adjustment for imaging confounders, the last one includes adjustment with rank inverse-normal transformation of all IDPs, age and neuroimaging confounders except for imaging centre. All sensitivity analyses were adjusted for twenty principal components of genetic ancestry.
